# Heat Waves Promote, but Cyclones Reverse, Coral-to-Seaweed Regime Shifts by Reshaping Reef Resilience

**DOI:** 10.64898/2026.09.27.754837

**Authors:** Russell J. Schmitt, Kai L. Kopecky, Andrew J. Brooks, Sally J. Holbrook

## Abstract

Disturbances alter ecosystem resilience by killing organisms and leaving behind material legacies. We tested how material legacies from marine heat waves and cyclones govern transitions between coral- and seaweed-rich states on tropical reefs. Heat waves kill coral tissue but leave their skeletons to erode gradually, whereas cyclones scour away emergent biota. Field experiments revealed that dead skeletons protect seaweeds from herbivory, catalyzing a transition to a seaweed-rich state that becomes self-sustaining after skeletons disappear. After four years, 73% of skeleton-added (‘heat wave’) plots transitioned to seaweeds compared to 0% of scoured (‘cyclone’) plots that lacked skeletons, while 87% of seaweed-rich plots scoured to simulate a cyclone stayed seaweed-free. Reef trajectories after natural disturbances were similar: four years after heat wave-induced coral mortality, 63% of time-series plots had transitioned to seaweeds with none recovering to coral; four years after a cyclone removed skeletons from a prior disturbance, none had transitioned to seaweeds and 60% had returned to coral. A disturbance not only can displace an ecosystem state, it can rapidly reshape the resilience landscape through the material legacies it creates or removes. Our findings also show that successive disturbances need not have cumulative negative effects: after a reef is driven into a seaweed-rich state, a cyclone can act as a rescue disturbance by removing established seaweeds and, when structural refugia are also removed, shrinking the seaweed stability domain. By reshaping resilience through contrasting material legacies, disturbance type and sequence can be as important as frequency and severity in determining ecosystem trajectories.

**Significance Statement:** Marine heat waves are becoming more frequent, and powerful cyclones stronger. Heat waves kill coral but leave their skeletons, whereas cyclones can cause comparable mortality but scour away skeletons and seaweeds. Decaying skeletons shelter seaweeds from herbivores, enabling seaweeds to proliferate, persist after skeletons disappear, and prevent coral recovery. Herbivores can better keep seaweeds suppressed on storm-scoured reefs, enabling coral to recolonize. Heat waves can promote shifts from coral to seaweeds, whereas cyclones can promote coral recovery. This rescue effect challenges the widespread view that successive disturbances are invariably harmful to coral or other ecosystems and shows that their effects depend not only on what they kill and leave behind, but also on how previous disturbances altered the ecosystem.

## Introduction

Climate change is reshaping ecosystems worldwide through the combined effects of changing drivers (e.g., rising mean temperatures) and acute disturbance events (e.g., storms, episodic heat waves) (1). Alterations in the frequency, severity, and spatial extent of pulse disturbances (1–3) are degrading ecosystem structure and functioning (4–6). The response of an ecosystem depends not only on the type of a disturbance, but also on its history of prior disturbances (5, 7, 8). Past disturbances can generate persistent material and information legacies - collectively known as ecological memory - that shape subsequent ecosystem dynamics (7–9). Material legacies are biotic or abiotic constituents left after a particular disturbance event (7, 10) such as standing dead trees in a forest after a drought (7) or standing skeletons of dead branching corals on a tropical reef after a severe bleaching event (11, 12). These legacies have been shown to modify post-disturbance trajectories (7, 9), and thereby have the potential to greatly affect resilience, that is, the system’s ability to absorb a perturbation and regain structure and function without transitioning to a self-sustaining alternative state (13).

In recent decades, many tropical reefs worldwide have undergone persistent regime shifts from coral- to seaweed-rich states, raising concern that changing disturbance regimes are eroding coral reef resilience (14–16). Historically, hydrodynamic forces from cyclonic storms were the dominant source of acute coral mortality (17). Although the global frequency of tropical cyclones is not consistently projected to increase with warming (18), the intensity of the strongest storms is expected to rise (19). More definitively, warming has driven rapid increases in the frequency and duration of marine heat waves (20), which cause mass coral bleaching mortality at unprecedented spatial scales and recurrence intervals (21). Consequently, severe marine heat waves now co-occur with powerful cyclones as major agents of disturbance on tropical reefs, representing a fundamental departure from the historical disturbance regime (6, 12).

Importantly, storms and heat waves leave behind qualitatively different material legacies on coral reefs (Fig. 1) (11, 12). Hydrodynamic forces from powerful storms detach coral colonies from the reef and grind down their calcium carbonate skeletons, leaving flatter, more open expanses of reef that may or may not have overburden patches of coral rubble (17, 22). Loose coral rubble can impair coral recovery by providing unstable settlement substrate (23), whereas sufficiently powerful hydrodynamic disturbance can scour emergent structure and expose stable primary reef (12) (Fig. 1B). By contrast, severe marine heat waves kill coral tissue but leave their standing skeletons attached to the reef (11, 12) (Fig. 1C), greatly expanding the availability of stable, structurally complex hard surfaces that can be colonized by seaweeds (upright macroalgae) and other sessile organisms. Herbivory by reef fishes is a key process promoting resilience of coral-rich states by suppressing seaweeds that compete with corals (15, 24). The slowly eroding dead skeletons can weaken this top-down control by providing physical refugia that reduce herbivore access to seaweeds (12, 25). When released from herbivory, seaweeds can proliferate and suppress coral growth, survival, and recruitment (26, 27), generating feedbacks that can reinforce seaweed-rich states (28, 29).

**Figure 1.**
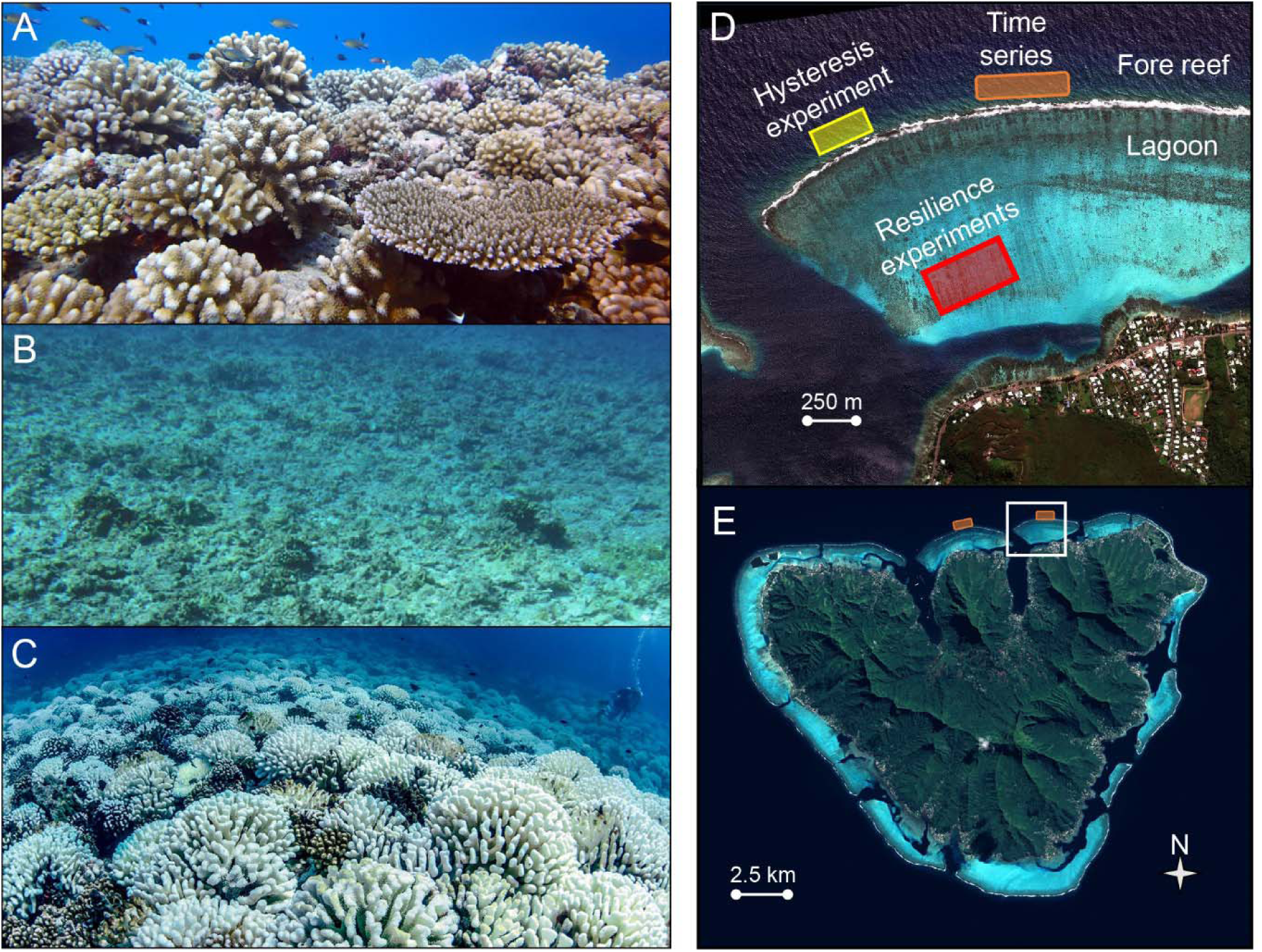
The fore reef of Moorea, French Polynesia at ∼ 10 m bottom depth (A) when live coral colonies covered ∼ 50% of the reef surface, (B) shortly after powerful storm waves removed coral colonies and other emergent biota and scoured the reef to primary attachment space, and (C) during a severe heat wave in 2019 that triggered mass coral bleaching mortality and left an extensive ghost forest of dead, gradually eroding coral skeletons. (D, E) Satellite images of Moorea, French Polynesia, showing the general experimental area on the north shore (D) in relation to the whole island (E). Locations of the experimental and time series sites are shown. NextView / WorldView satellite imagery (Copyright 2011, DigitalGlobe, Inc.) was provided under a National Geospatial-Intelligence Agency NextView License agreement with the MCR LTER for NSF-sponsored MCR research purposes only, which permits posting of reduced-resolution imagery devoid of metadata. Photo credits: (A) Russell J. Schmitt, (B) Kai L. Kopecky, (C) Peter J. Edmunds

Here we use *in situ* hysteresis experiments (tests for path dependence) and resilience experiments (tests for non-recovery) to determine how the contrasting material legacies from heat waves and cyclonic storms reshape regime-shift pathways and coral-seaweed dynamics. To test whether disturbance legacies alter the resilience landscape, we repeated on the same fore reef tract a hysteresis experiment conducted after a Category 4 cyclone in 2010 (29), this time following a severe marine heat wave that occurred in 2019 (Fig. 1D). The cyclone not only scoured away most emergent biota on the fore reef but also removed remnants of a ghost forest of dead coral skeletons left by a preceding boom-and-bust outbreak of a coral predator (30–32). By contrast, standing dead skeletons left by the 2019 heat wave eroded gradually over several years. We used results of these hysteresis experiments to construct a conceptual resilience landscape that distinguishes how disturbances can simultaneously reposition ecosystem state on the landscape by changing the abundance of organisms and also reshape the landscape itself by altering physical habitat and the ecological interactions it mediates. Our resilience experiments, grounded by the two disturbance events, revealed transition pathways and the likelihood of coral-to-seaweed regime shifts. We then tested whether the experimentally predicted trajectories emerged on unmanipulated reefs using time series of coral and seaweed abundances following the two natural disturbance events. Our resilience-landscape framework links contrasting disturbance legacies to transitions among alternative states in ecosystems structured by habitat-forming foundation species.

## Results

### Effect of a Dead Skeleton Ghost Forest on Hysteresis in the Herbivore-Seaweed Relationship

Hysteresis in driver–response relationships—when transitions between stability domains occur at different values (thresholds) of the driver, depending on the direction of the transition—creates a range of the driver values under which alternative domains are possible. Our Hysteresis Experiments (tests of path dependence) addressed whether and how the driver-response relationship between herbivorous fish and seaweeds differed depending on the starting assemblage and extent of refugia provided by dead coral skeletons in the environment. The initial assemblages represented two endpoints of algal succession on coral reefs: thin epilithic turf that initially colonizes newly opened hard surfaces and can subsequently be colonized by coral or seaweeds, and a late-successional assemblage of fleshy brown seaweeds (primarily *Turbinaria ornata*) that strongly inhibits coral colonization. These two initial algal states were subjected to the same experimental gradient in herbivory (from none to ambient) to estimate the driver-response relationship. The hysteresis experiments were done across three different structural contexts where the presence of dead coral skeletons varied within experimental plots and across the surrounding reefscape. The results revealed that herbivore thresholds for suppressing seaweed proliferation versus removing established seaweeds varied greatly depending on physical refugia provided by dead skeletons (Fig. 2).

**Figure 2.**
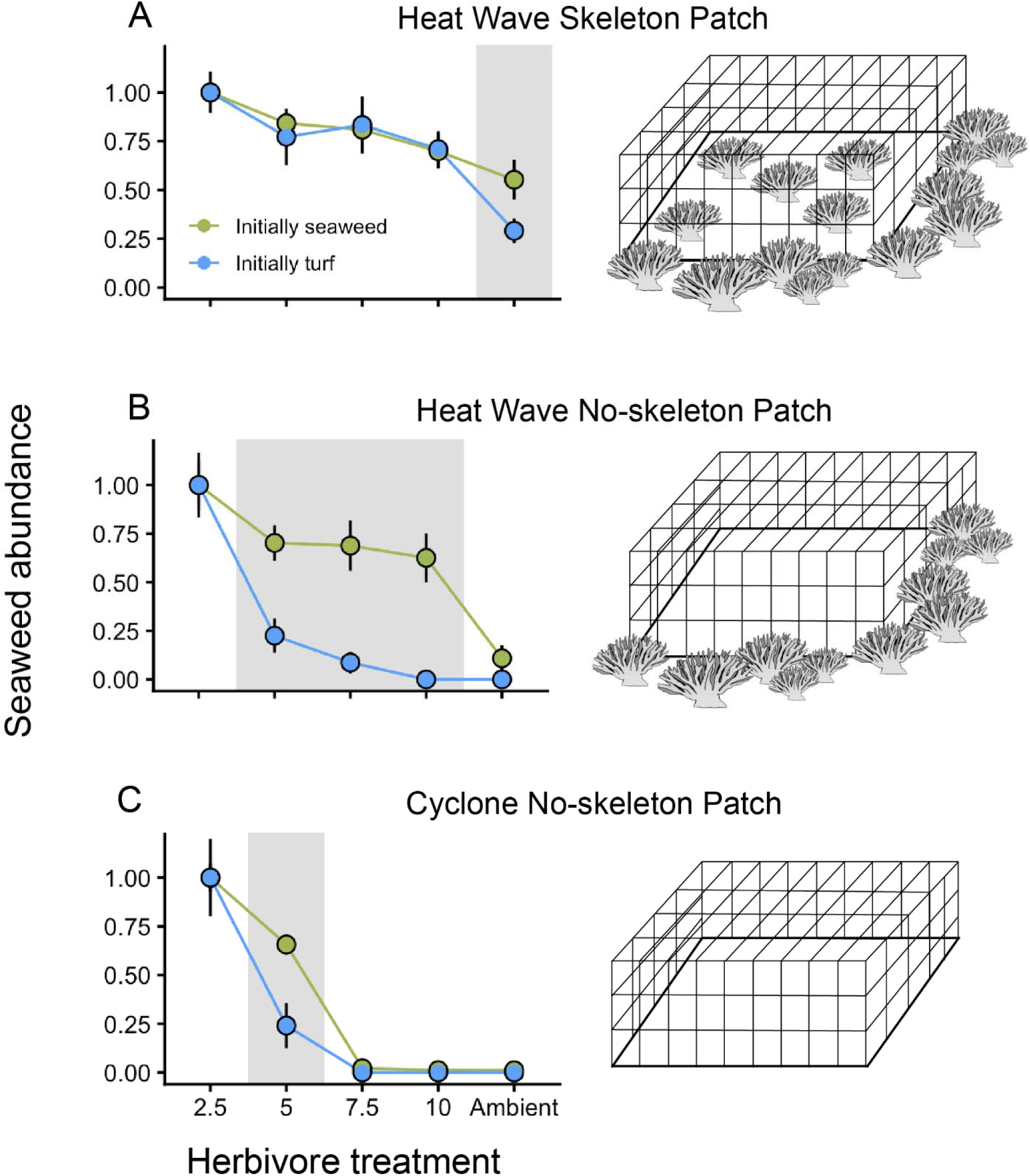
Results of experimental tests for hysteresis in the relationship between seaweeds and herbivores as a function of the amount of algal refugia provided by dead coral skeletons within an experimental patch and in the surrounding reefscape. Experiments were conducted at the same fore reef location after two major disturbance events: a severe marine heat wave in 2019 (A & B) and a Category 4 cyclone in 2010 (C); data for (C) replotted from (29). Panels are ordered from the greatest (A) to least (C) amount of protection from herbivory provided by dead coral skeletons within and surrounding an experimental patch. In each case, an herbivore exclusion cage is shown to illustrate the structural context of the reef environment and experimental patch. Graphs are the herbivore-seaweed relationships as a function of initial algal state (i.e., forward and reverse driver-response trajectories) for (A) a plot that contained dead skeletons similar to the surrounding reef immediately after the 2019 marine heat wave (Heat Wave Skeleton Patch treatment), (B) an open patch embedded within the post-heat wave forest of dead skeletons (Heat Wave No-skeleton Patch treatment), and (C) an open, structure-free patch embedded within the post-cyclone reefscape of exposed primary substrate (Cyclone No-skeleton Patch treatment). Data are the mean (± SE) final cover of seaweeds, standardized to the mean of the lowest (2.5) herbivory treatment, for two initial algal states: a mature stand of seaweeds (green circles); and thin epilithic turf (blue circles) that is the first colonizer of denuded hard substrates and can be colonized by either coral or seaweeds. Gray polygons denote the range of herbivore abundance over which bistability between seaweed and coral–epilithic turf states may occur, based on statistically significant differences between the two initial states at a given herbivory level.

When skeletons were present within a patch as well as the surrounding reefscape (Heat Wave Skeleton Patch treatment), seaweeds became dominant across most of the experimental herbivory gradient, with little evidence that initial algal state (turf vs. mature seaweed) altered the herbivore– seaweed relationship (Fig. 2A; for non-normalized trajectories, see Fig. S1). Initial algal state did not significantly affect seaweed percent cover across the herbivory gradient (Table S1; Herbivore × Initial State Interaction: F_4,50_ = 1.1; *P* = 0.39; Fig. S1A), although a significant interaction was detected for seaweed biomass (Table S1; F_4,50_ = 3.8; *P* < 0.01; Fig. S1B). Normalized trajectories suggested that any region of potential bistability was confined to the ambient-herbivore treatment (gray region in Fig. 2A), where normalized seaweed cover differed significantly between initial states (Student’s t test: t_8_ = 2.8; *P* = 0.02; arcsine-transformed data). Even at ambient herbivory, however, both putative alternative algal states were characterized by high mean seaweed abundance (Figs. 2A, S1A). Thus, when dead skeletons were present both within experimental plots and throughout the surrounding reefscape, seaweed dominance occurred across nearly the entire experimental gradient in herbivory, with little evidence for distinct high- and very low-seaweed states.

By contrast, when an open patch that lacked dead skeletons was embedded in a ghost forest of dead skeletons (Heat Wave No-skeleton Patch treatment), initial-state trajectories remained widely separated in seaweed abundance across much of the herbivory gradient, indicating a broad region of potential bistability in which herbivory sufficient to prevent seaweed establishment was insufficient to reverse an established seaweed state (gray region in Fig. 2B). The herbivore–seaweed relationship differed significantly between initial algal states for both seaweed percent cover (Table S1; Herbivore x Initial State Interaction: F_4,50_ = 4.8; *P* < 0.001; Fig. S1C) and biomass (F_4,50_ = 3.2; *P* < 0.005; Fig. S1D). These interactions resulted from divergence between the initial-state trajectories as herbivore levels increased (Fig. S2): established seaweeds persisted at levels of herbivory sufficient to prevent seaweed proliferation from turf in all treatments except ambient herbivory (Fig. 2B).

When an open patch that lacked dead skeletons was embedded on the flattened, skeleton-free reefscape after Cyclone Oli (Cyclone No-skeleton Patch), the herbivore–seaweed relationship showed statistical evidence of hysteresis at only one herbivory treatment, well below the prevailing abundance of herbivores at the site (Fig. 2C; 29). Consequently, the region of potential bistability was substantially narrower in the skeleton-free post-cyclone reefscape than in the post-heat-wave reefscape.

### Effect of Disturbance Type on State Transition Probabilities

To experimentally evaluate whether (a) marine heat waves can catalyze a critical transition from a coral- to seaweed-rich regime and (b) a powerful cyclone can reverse that regime shift, we concurrently carried out two 4-year-long Resilience Experiments (tests of non-recovery) (33): the Coral Resilience Experiment and the Seaweed Resilience Experiment. Replicates of the two experiments were spatially interspersed among 60 adjacent lagoon patch reefs (Fig. 1D) where seaweed-rich and turf-coral states have been shown to behave dynamically as alternative stability domains (29, 34). We classified replicate plots as seaweed-rich when seaweed cover exceeded 25.0%, the first quartile observed on unmanipulated patch reefs where seaweeds had persisted as major emergent space holders for the preceding decade (Fig. S3A) (29, 34). This threshold is ecologically meaningful because seaweed cover at this level inhibits coral recolonization on these patch reefs (34). The brown algae *Turbinaria ornata* and *Sargassum pacificum* together accounted for > 95% of seaweed cover and biomass on these unmanipulated patch reefs.

#### Coral Resilience Experiment

Initially, all patch reefs used in the Coral Resilience Experiment were devoid of seaweeds (Fig. 3A), cleared of other emergent biota and covered by closely cropped epilithic turf that can be colonized by either coral or seaweeds. At the start of the experiment, dead *Pocillopora* skeletons were affixed to half of the plots (Coral Heat Wave treatment; Fig. 3B) in a manner that mirrored the spacing of dead branching corals following the 2019 heat wave event (12). The remaining turf-covered, skeleton-free plots (Coral Cyclone treatment) represented the structural condition of the reef surface following a powerful storm such as Cyclone Oli in 2010, which scours away corals, seaweeds, and other emergent biota. After 24 months, when approximately 60% of the experimental skeletal material remained (Fig. S4), 67% of the 15 Coral Heat Wave replicates had transitioned to a seaweed-rich state (Fig. 3D, E), whereas none of the Coral Cyclone replicates had done so (Fig. 3F; Fisher’s exact test, *P* < 0.0002). The two principal species of seaweeds that colonized the Coral Heat Wave plots were *Turbinaria ornata* and *Sargassum pacificum* (Figs. 3E, H).

**Figure 3.**
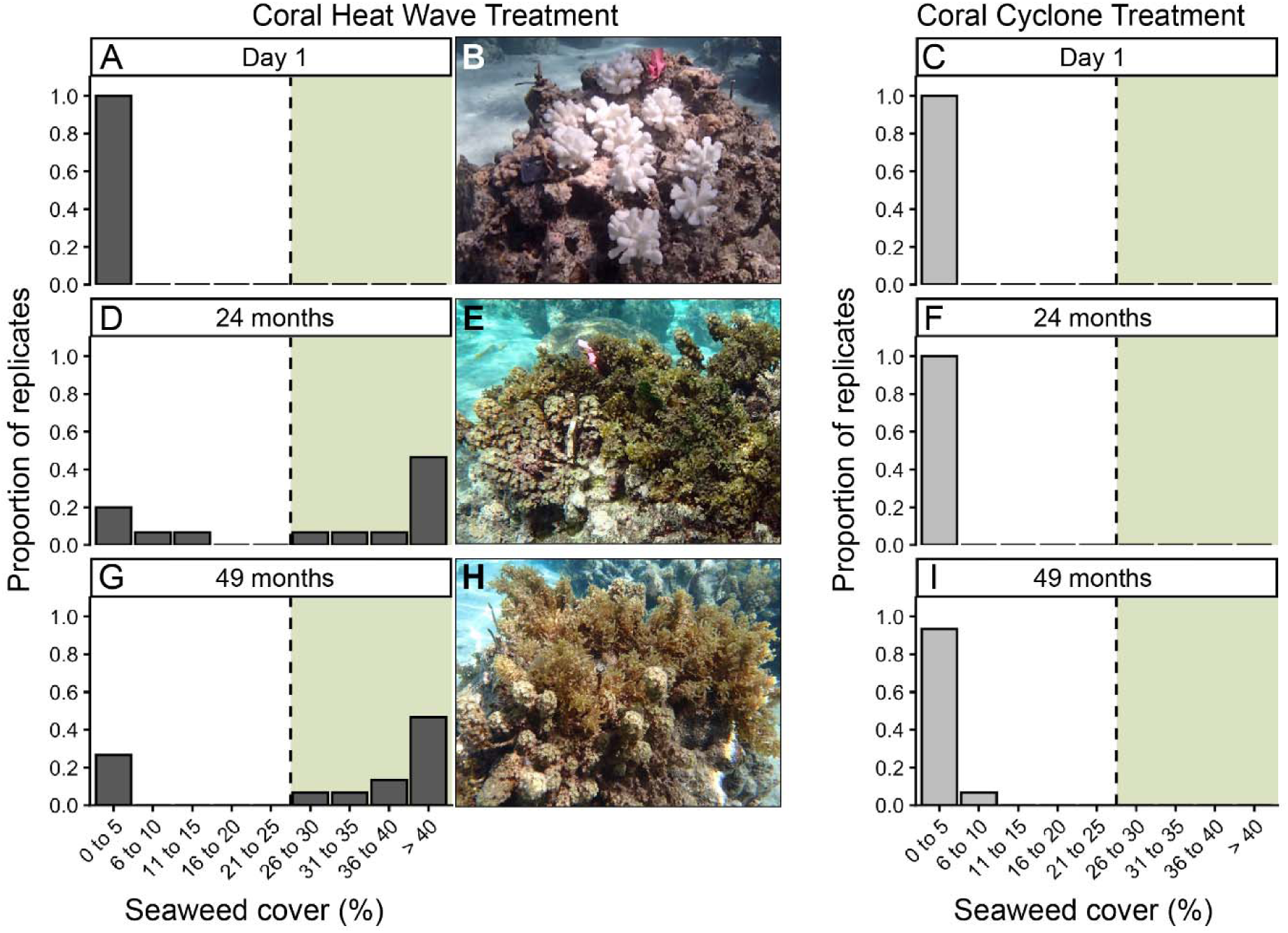
Results of the Coral Resilience Experiment, which tested whether the material legacy of a marine heat wave—gradually eroding dead coral skeletons—could facilitate a coral-to-seaweed regime shift. Graphed are frequency distributions of seaweed cover for the Coral Heat Wave treatment (A, D, G) and the Coral Cyclone treatment that received no dead coral skeletons (C, F, I). The vertical dashed line on each histogram denotes the 25% seaweed-cover threshold used to distinguish seaweed-depauperate (white) from seaweed-rich (green shading) states. Distributions are shown at the start of the experiment (A, C), after 24 months, when slightly more than 60% of the experimental skeletal material remained (D, F), and after 49 months, four months after complete disappearance of the skeletons had been confirmed (G, I). Images (B, E, H) show the same Coral Heat Wave replicate at the three sampling times; *Turbinaria ornata* and *Sargassum pacificum* were the principal seaweeds that colonized and persisted on Coral Heat Wave reefs (E, H). Photo credits: R. J. Schmitt.

Our estimated erosion rate indicated that experimentally added skeletons would disappear by 42 months, and direct measurements at 45 months confirmed their complete disappearance (Fig. S4). Because the turnover time of adult *Turbinaria* is roughly 4 months in our system (29, 35), we assessed state persistence at 49 months, at least one complete turnover time after the skeletal refugia had disappeared. At 49 months, 73% of Coral Heat Wave replicates were seaweed-rich (Figs. 3G, H), including every replicate that had transitioned by 24 months, whereas none of the Coral Cyclone replicates had transitioned to seaweeds (Fig. 3I; Fisher’s exact test, *P* < 0.0001). For the Coral Heat Wave replicates that transitioned to seaweeds by 49 months, the mean (± 1 SE) percent cover of seaweeds was 49.6% (± 6.2%), whereas that on the Coral Cyclone plots was 4.0% (± 1.1%) (Welch’s t-test: t_13_ = 9.25; *P* < 0.0001). Mean seaweed cover among Coral Heat Wave replicates in the seaweed-rich state did not differ between 24 months (55.4% ± 6.5%) and 49 months (49.6% ± 6.2%; Welch’s t test: t_19_ = 0.65; *P* > 0.53), indicating that complete erosion of the skeletal refugia produced no detectable thinning of established seaweed cover. A transient pulse of dead coral skeletons catalyzed a transition to high seaweed cover that, once established, persisted after the skeletal legacy disappeared and the adult seaweed population turned over.

#### Seaweed Resilience Experiment

To test whether storm-driven removal of established seaweeds could reverse the seaweed-rich state, we removed emergent seaweeds while leaving the underlying structural heterogeneity of the reef surface—and hence potential physical refugia for seaweeds— unchanged. Before manipulation, all Seaweed Cyclone and unmanipulated Seaweed Control plots were seaweed-rich and dominated by *Turbinaria* with some *Sargassum*, with similar initial seaweed cover (mean ± SE: Seaweed Cyclone treatment, 50.3 ± 3.3%; Seaweed Control, 56.3 ± 1.8%; Welch’s t test: t_22_ = 1.6, *P* > 0.13). After 49 months, 87% of Seaweed Cyclone replicates had failed to recover to the seaweed-rich state (Figs. 4D, E), whereas all Seaweed Control replicates remained seaweed-rich (Fig. 4F; Fisher’s exact test, *P* < 0.0001). Seaweed Cyclone plots that failed to recover had only 4.6% (± 1.2%) seaweed cover at 49 months, indistinguishable from the 4.0% (± 1.1%) observed in Coral Cyclone plots that had begun the experiment in the turf state (Welch’s t test: t_25_ = 0.37; *P* = 0.72). Likewise, the probability of remaining seaweed-depauperate did not differ between the two treatments (Fisher’s exact test, *P* = 0.48). Removal of established seaweeds shifted most plots to a seaweed-depauperate, coral-invasible turf state that persisted for at least 4 years.

**Figure 4.**
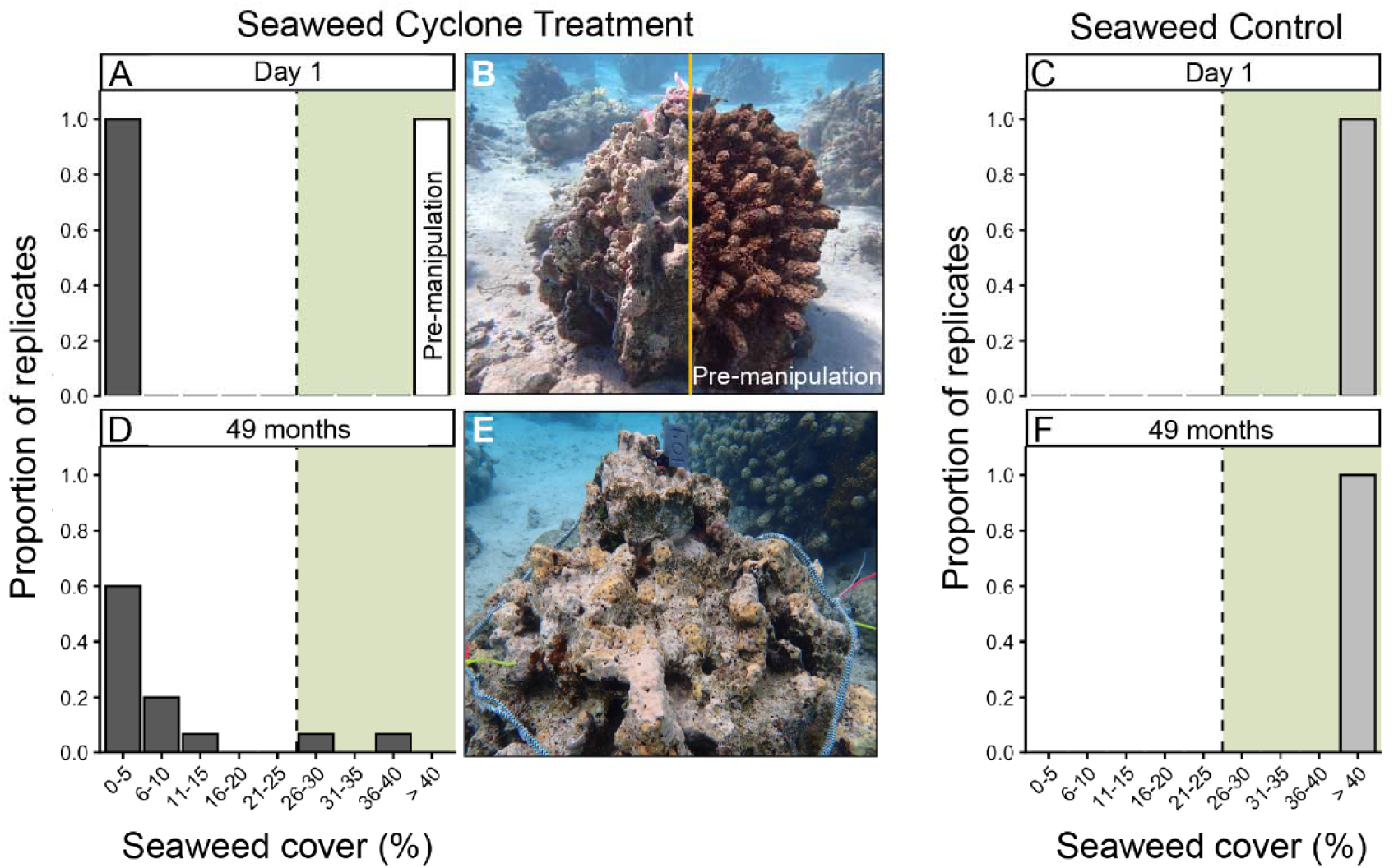
Results of the Seaweed Resilience Experiment, which tested whether storm-like removal of established seaweeds could reverse the seaweed-rich state without altering physical refugia or otherwise reshaping the resilience landscape. Graphed are frequency distributions of seaweed cover for the Seaweed Cyclone (A, D) and unmanipulated Seaweed Control treatments (C, F) at the start of the experiment (A, C) and after 49 months (D, F). The vertical dashed line denotes the 25% seaweed-cover threshold used to distinguish seaweed-depauperate (white) from seaweed-rich (green shading) states. The split image in (B) shows the same Seaweed Cyclone replicate immediately before (right) and after (left) removal of established seaweeds; (E) shows that replicate at the 49-month endpoint. Photo credits: R. J. Schmitt.

### Natural Post-disturbance Transition Probabilities and Dynamics

To classify natural fore reef plots as coral-rich or coral-depauperate, we used the first quartile of coral cover (25.2%; median = 35.3%) measured in 2006, 15 years after the previous major coral-killing disturbance (Fig. S3). This threshold was nearly identical to the 25.0% threshold used to distinguish seaweed-rich from seaweed-depauperate states. Using these thresholds, only 1 of 351 plot-years (0.3%) was simultaneously coral-rich (≥ 25.2% cover) and seaweed-rich (≥ 25.0%), reflecting the strong negative covariation between these competing space holders on the fore reef. Thus, coral-rich and seaweed-rich conditions were nearly mutually exclusive, providing an objective basis for quantifying post-disturbance transitions between these contrasting reef states.

#### Four-year State Transition Probabilities

At the onset of the severe marine heat wave in 2019, all unmanipulated fore reef plots (Fig. 1D) were coral-rich and seaweed-depauperate (Fig. 5C). The heat wave caused mass bleaching mortality that initially killed ∼ 80% of the coral present (Fig. 5C), leaving a widespread ghost forest of dead skeletons composed largely of branching corals in the genus *Pocillopora* (Fig. 1C) that gradually eroded for years after the heat wave. Four years after the heat wave—the same timescale over which resilience was experimentally assessed—63% of plots had transitioned to the seaweed-rich state (Fig. 5A), whereas none had recovered to the coral-rich state (Fig. 5B). Coral cover remained extremely low (≤ 5%) on 63% of plots (Fig. 5B).

**Figure 5.**
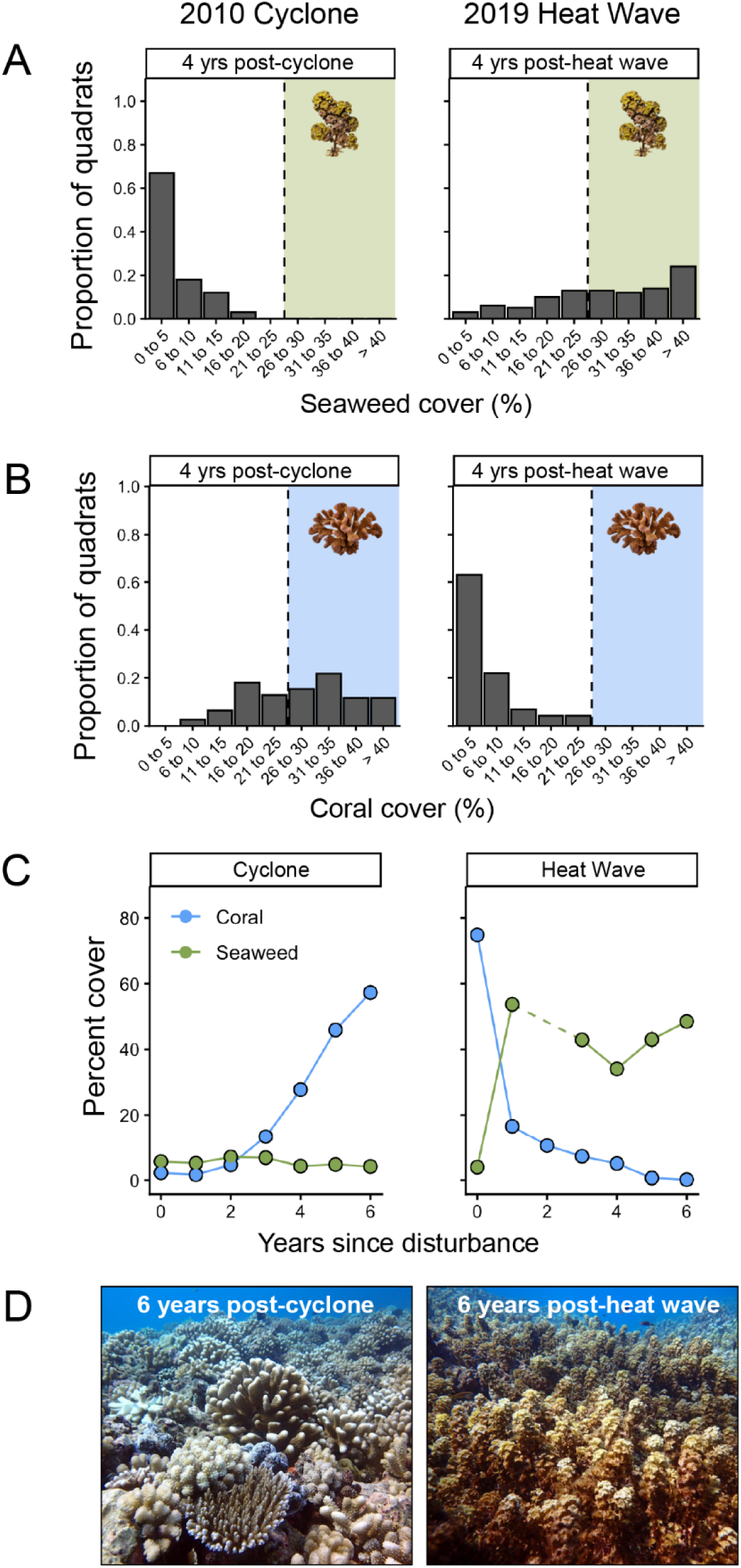
Natural post-disturbance dynamics of seaweeds and coral on the north shore fore reef of Moorea at 10 m depth following the Category 4 Cyclone Oli in 2010 (left column) and the severe marine heat wave in 2019 (right column). (A) Frequency distributions of seaweed cover four years after each disturbance, based on 100 quadrats following Cyclone Oli and 100 quadrats following the heat wave. (B) Frequency distributions of live coral cover four years after each disturbance, based on 78 permanent quadrats following Cyclone Oli and 73 quadrats following the heat wave. The vertical dashed lines in (A) and (B) denote the independently estimated thresholds used to classify quadrats as seaweed-rich (green shading) or coral-rich (blue shading). (C) Trajectories of mean seaweed cover (green circles) and live coral cover (blue circles) during the six years following each disturbance. (D) The fore reef site of the Hysteresis Experiments six years after Cyclone Oli (left) and six years after the 2019 heat wave (right). Six years after the cyclone, live coral had recovered to become the predominant space holder at our experimental site, whereas six years after the heat wave, the reef had transitioned to a seaweed-rich state. Photo credits: R. J. Schmitt.

Three years before Cyclone Oli, live coral occupied ∼ 50% of attachment space on the fore reef, but a boom-and-bust outbreak of the voracious coral predator crown-of-thorns seastar (*Acanthaster planci*) reduced cover of living coral tissue to ∼1% before the cyclone struck (30, 32). Cyclone Oli then swept away the forest of dead coral skeletons left by the crown-of-thorns seastar outbreak (Fig. 1B), leaving a flattened, rubble-free reefscape with extremely low cover of both coral and seaweeds (36) (Fig. 5C). Four years after Cyclone Oli, no plot had transitioned to the seaweed-rich state (Fig. 5A), whereas 60% had recovered to the coral-rich state (Fig. 5B). Consequently, the four-year probability of transition to the seaweed-rich state was substantially lower following Cyclone Oli than following the 2019 heat wave (χ²_₁_= 89.1, *P* < 0.0001), whereas the probability of recovery to the coral-rich state was substantially higher (χ²_₁_ = 61.1, *P* < 0.0001). Thus, the two disturbances generated sharply different state transition probabilities over the same four-year timescale despite leaving similarly low coral and seaweed abundances immediately afterwards.

### Post-disturbance Population Trajectories

These contrasting state-transition probabilities arose from opposing changes in coral and seaweed abundance following the two types of disturbance (Fig. 5C). After the 2019 heat wave, coral cover declined from 72% to 16% in the first year, while seaweed cover increased from < 5% to > 50% (Fig. 5C). Seaweeds rapidly colonized spaces beneath and between the branches of dead coral skeletons and initially consisted primarily of ruffled and decumbent forms of the brown alga *Lobophora*. Over the next 3 years, coral cover declined further, primarily because of competitive encroachment by *Lobophora* (25), while seaweed cover remained high at ∼ 35–50% (Fig. 5C). No strong hydrodynamic event removed the remnant dead skeletons during the 6 years following the heat wave, and they instead gradually eroded throughout this period. As erosion exposed primary attachment space, it was increasingly colonized by *Turbinaria ornata* (Fig. 5D).

Following Cyclone Oli, which scoured away the dead coral skeletons produced by the preceding crown-of-thorns seastar outbreak, seaweed cover remained < 8% throughout the first 6 years after the storm (Fig. 5C). Intense herbivory by reef fishes suppressed seaweeds and prevented them from capturing newly available space (31, 36, 37), leaving exposed primary attachment space available for colonization by sexually produced coral propagules, likely originating from populations that escaped the severe fore reef disturbance (38). Coral cover increased steadily following the cyclone, returning to its pre-disturbance level of ∼ 50% within 6 years (Figs. 5C, D) (32).

### Graphical Resilience Landscape Model and Implications for Disturbance-induced Regime Shifts

We extended our previous conceptual model of how the contrasting material legacies of marine heat waves and cyclonic storms alter the herbivore-seaweed relationship and thereby reshape the resilience landscape (11). Figure 6A depicts the revised conceptual landscape, informed by the experimentally observed herbivore-seaweed relationships in Figure 2, which represent three positions along a gradient of physical refugia for seaweeds. This model illustrates how seaweed abundance and the potential for alternative states could vary jointly with herbivore abundance and the availability of physical refugia from herbivory. Colored cross-sections through the surface (brown, A; orange, B; blue, C) correspond to the empirical herbivore–seaweed relationships in Figs. 2 A–C, respectively, spanning maximal to minimal refugia. The correspondingly colored spheres indicate the inferred ecosystem states at a common herbivore abundance across the refugia gradient, approximately corresponding to the long-term ambient abundance of herbivores at the study site (black circle on herbivore axis). Thus, changing physical refugia could shift ecosystem state at a given herbivore abundance. The response surface shows how the regions supporting coral dominance, seaweed dominance, or bistability—and hence the resilience of the alternative states—shift with herbivore abundance and physical refugia. The positions of the severe heat wave 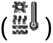 and cyclonic storm 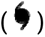 icons along the refugia axis illustrate how their contrasting material legacies can move the system in opposite directions along this axis, thereby reshaping the resilience landscape without necessarily altering herbivore abundance.

**Figure 6.**
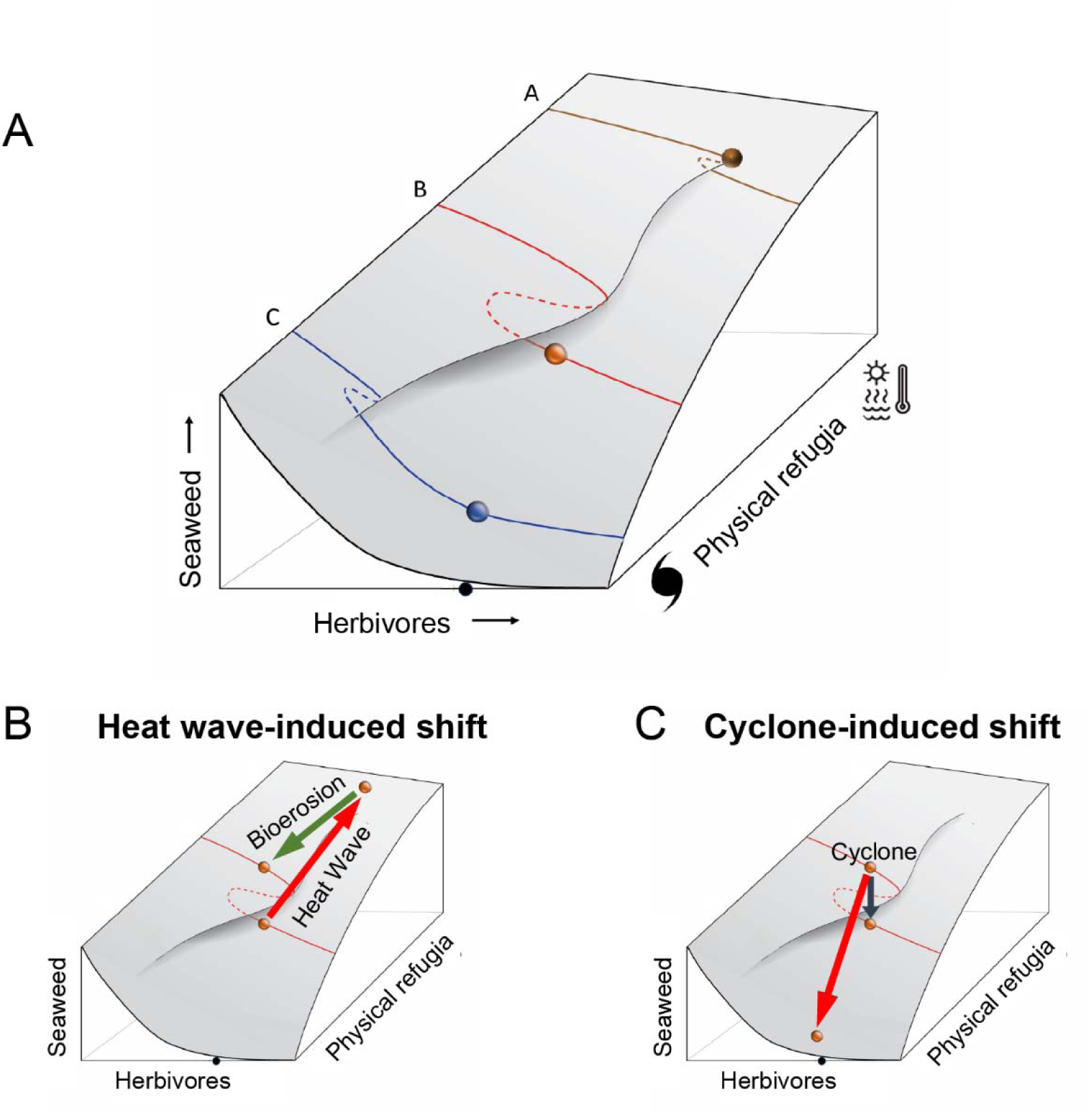
(A) Conceptual resilience landscape (multi-driver response surface) for our system, informed by results of the Hysteresis Experiments (Fig. 2), depicting the equilibrium abundance of seaweeds as a joint function of herbivore abundance and physical refugia that reduce per capita herbivore effectiveness. Colored lines A, B, and C represent cross-sections through the resilience landscape at three positions along the algal-refugia gradient corresponding to the hysteretic trajectories in Fig. 2A (maximal refugia), 2B, and 2C (minimal refugia), respectively. Colored spheres represent the inferred ecosystem states at the same ambient herbivore abundance (black dot on herbivore axis) across the refugia gradient. The relative positions of the severe marine heat wave 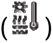 and cyclonic storm 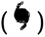 icons along the refugia axis indicate that these disturbance types tend to alter physical refugia in opposite directions, thereby rapidly reshaping the resilience landscape without directly altering herbivore abundance. (B) Hypothesized pathway by which a severe marine heat wave can trigger a coral-to-seaweed regime shift without altering herbivore abundance. The bivariate physical refugia–seaweed diagram holds herbivore abundance constant across the refugia gradient. Initially, the pre-heat-wave ecosystem (orange sphere) lies within the region of coral–seaweed bistability, where both states are stable. By killing coral but leaving dead skeletons in place, a severe marine heat wave acts through two pathways: coral mortality displaces the ecosystem toward the seaweed-rich state, while the resulting ghost forest reduces per capita herbivore effectiveness and expands the stability domain of the seaweed-rich state. Together, these effects can carry the ecosystem across a stability boundary into the seaweed-rich state (red arrow). As dead skeletons gradually erode and structural refugia decline, the resilience landscape shifts back toward its prior configuration (green arrow). Due to hysteresis, however, the ecosystem can remain in the seaweed-rich state rather than retracing the trajectory by which it formed. (C) Hypothesized pathways by which a cyclonic storm can shift an ecosystem out of the seaweed-rich state. Within the region of coral–seaweed bistability, removal of established seaweeds can displace the ecosystem across a stability boundary toward the coral-rich state without changing the resilience landscape (black arrow). When a cyclone also removes structural refugia, it increases per capita herbivore effectiveness and contracts the stability domain of the seaweed-rich state, reshaping the resilience landscape in the same direction (red arrow).

## Discussion

The ecological consequences of a disturbance depend not only on the magnitude of mortality and state of the system but also on the material legacies that remain after organisms are killed (7–12). Disturbance type matters because material legacies differ (11, 12). Our experiments revealed that heat waves and cyclones can have opposing effects on coral reef resilience: the heat-wave material legacy (dead skeletons) degraded the resilience of the coral-rich state, whereas a cyclone legacy (reduced physical structure) degraded the resilience of the seaweed-rich state. Consistent with these opposing experimental effects, the fore reef benthic community of Moorea followed sharply divergent trajectories after two major disturbances that caused similarly severe coral loss but left contrasting material legacies. The 2019 marine heat wave left a slowly eroding ghost forest of dead coral skeletons that provided physical refugia for seaweeds and reduced their exposure to herbivory (12). By contrast, Cyclone Oli removed a forest of dead skeletons that had just been created by a boom-and-bust outbreak of a coral predator, leaving a flattened, open, rubble-free reefscape where seaweeds thereafter remained sparse and coral subsequently recovered (31, 32, 36, 39). Following Cyclone Oli, grazing pressure was demonstrated to be sufficient to prevent young life stages of seaweeds from proliferating into adults (31, 36, 40). This clearly was not the case following the 2019 heat wave as seaweeds rapidly proliferated, consistent with experimental evidence that skeletons hinder grazer and browser access to algal resources (12). Our experiments isolated the mechanism, showing that even temporary physical refugia for algae reduce the effectiveness of a given level of herbivory, thereby increasing the probability that a disturbed reef transitions to a seaweed-rich state.

Material legacies can modify processes that stabilize a state (7, 12). Our hysteresis experiments show how changes in the effectiveness of herbivory can reshape the resilience landscape by altering the stability domains of coral and seaweed states. Herbivory is a major stabilizing process on coral reefs because it suppresses seaweeds that compete with corals (15, 16, 24, 28). Kopecky et al. (11) proposed that disturbance-generated physical structure can modify this stabilizing process: dead coral skeletons provide refugia from herbivory, expanding the seaweed stability domain and the range of herbivore abundance over which coral and seaweeds can be bistable. Our hysteresis experiments provided empirical support for this theoretical prediction across three contrasting structural contexts. The bistability domain occupied only a narrow portion of the herbivory gradient when a cyclone had scoured away emergent biota and reduced physical structure, leaving an open, flattened reefscape (Cyclone No-skeleton Patch experiment) (29). For a patch without physical structure in a ghost forest of dead skeletons (Heat Wave No-skeleton Patch treatment), already established seaweeds persisted across nearly the entire herbivory gradient, including levels of herbivory sufficient to prevent young seaweeds from escaping control and reaching maturity. This greatly broadened the region of potential bistability. However, when the patch within the ghost forest also contained dead skeletons (Heat Wave Skeleton Patch treatment), the seaweed-rich state occurred across the entire herbivory gradient. Independent evidence supports this mechanism: branching coral structure and small-scale structural refugia reduce herbivore access to seaweeds (41, 42), dead coral skeletons greatly reduce grazing and browsing by herbivorous fishes following bleaching (12), and experimental retention of those skeletons promotes seaweed proliferation (25). Disturbance-generated refugia can reshape the resilience landscape by shifting the boundaries and relative extent of coral, seaweed, and bistable domains (Fig. 6).

The ecological effects of the dead-skeleton legacy outlasted the skeletons themselves. In our resilience experiments, every Coral Heat Wave replicate that had transitioned to the seaweed-rich state in the presence of skeletons remained in that state well after the skeletal refugia had completely eroded, with seaweed cover remaining high despite turnover of adult plants. Conversely, most Seaweed Cyclone plots in which established seaweeds were removed remained seaweed-depauperate throughout the four-year experiment; the closely cropped turf state in these replicates has been shown to be readily colonized by corals at this site (34). Together with the hysteresis experiments, these results demonstrate strong path dependence, consistent with evidence that disturbance legacies can alter ecosystem resilience and recovery trajectories in other ecosystems (7, 43): levels of herbivory sufficient to prevent young seaweeds from reaching maturity need not be sufficient to reverse seaweed dominance once mature populations have developed (29, 34). The life history of *Turbinaria ornata*, the dominant seaweed in our system, illustrates a mechanism by which the seaweed-rich state can outlast the skeletal legacy that facilitated its development. Young *T. ornata* are highly vulnerable to herbivorous fishes, whereas vulnerability declines sharply with size (44, 45), and established, heavily-defended adults provide associational refuge for vulnerable recruits (44). A stage-structured model showed that maturation from herbivore-vulnerable seaweed recruits to resistant adults can generate alternative coral- and seaweed-dominated states over a broad range of herbivory (46). Because this type of stage structure is common in biological populations, the mechanism may be broadly applicable (46). Established *T. ornata* can persist under levels of herbivory sufficient to eliminate seaweeds that remain more palatable (47). Nutrient enrichment can further promote escape from herbivory by accelerating growth through the highly vulnerable recruit stage and speeding the development of structural defenses against herbivores (45). Dead skeletons can provide a temporary window for vulnerable seaweeds to escape herbivory and establish (12, 25); once populations contain herbivore-resistant, reproductive adults, their persistence and replenishment need not depend on the skeletal refuge that enabled establishment. Thus, the ghost forest can act as a transient catalyst for a state change that becomes self-sustaining: as the skeletons erode and physical refugia decline, the system can remain seaweed-rich rather than retracing the trajectory by which it formed (Fig. 6C).

The contrasting effects of heat waves and cyclones reveal that disturbances can simultaneously reposition an ecosystem within its resilience landscape and reshape the landscape itself (Fig. 6). Previous stability-landscape models distinguish changes in environmental conditions that alter the configuration of the landscape from pulse disturbances that displace an ecosystem within it (48–51). Pulse disturbances can themselves rapidly reshape the landscape through the material legacies they create or remove. A severe marine heat wave reduces coral abundance, displacing the ecosystem toward the seaweed-rich state, while the resulting ghost forest reduces per capita herbivore effectiveness and expands the stability domain of the seaweed-rich state. A powerful cyclone can act through the same two pathways in the opposite direction. Removal of established seaweeds alone can displace the ecosystem far enough toward the coral-rich state to cross a stability boundary without changing the resilience landscape (Figs. 4 and 6C), whereas removal of structural refugia increases per capita herbivore effectiveness and contracts the stability domain of the seaweed-rich state. These effects can drive the ecosystem toward the coral-rich state while reshaping the resilience landscape in the same direction (Fig. 6C). Conversely, a sufficiently large increase in refugia following a heat wave could expand the seaweed stability domain until it encompasses the existing reef state, without any reduction in herbivore abundance (Fig. 6B) (11).

Because different disturbance types can drive ecosystem trajectories in opposing directions, successive major disturbances need not have cumulative negative effects. Following a coral mass-mortality event that leaves a skeleton ghost forest, a subsequent cyclone can remove established seaweeds and structural refugia provided by both dead coral skeletons and the reef matrix. Removal of established seaweeds disrupts the feedbacks that stabilize the seaweed-rich state, while removal of structural refugia reduces the probability that vulnerable life stages will escape herbivory (Fig. 6C). The post-Cyclone Oli trajectory illustrates this reversal: removal of dead skeletons was followed by persistently sparse seaweeds and rapid coral recovery (Fig. 5) (31, 32, 36, 39). Consistent with this pattern, coral-rich reefs severely damaged by a Category 5 cyclone in Fiji shifted to turf rather than seaweeds, rapidly returning to a coral-rich state over the following four years (52). These examples point to a counterintuitive role for disturbance: a powerful cyclone can act as a rescue disturbance for a reef driven toward or into a seaweed-rich state by an earlier disturbance. Whether coral recovery follows such a rescue disturbance remains context dependent because coral establishment following storms requires sufficient abundance of key herbivore functional groups and coral recruitment and can be impeded by severe damage to the reef matrix or extensive patches of unstable rubble (23, 31, 32).

As climate change continues to alter disturbance regimes, the goal of understanding their ecological consequences (1–10) will require considering disturbance type and sequence as well as intensity and frequency (53). Disturbances that cause similar mortality can leave different material legacies, while the consequences of a subsequent disturbance can depend on the state (54) and legacy created by those that preceded it. Material legacies provide a mechanism of ecological memory through which disturbance history influences subsequent ecosystem dynamics (7–9), a phenomenon documented for the remains of foundation species across terrestrial and marine ecosystems (9). Because the effects can persist after the physical legacy itself disappears, a transient legacy can leave a persistent ecological imprint—a “ghost of legacies past” (55). Predicting ecosystem resilience under changing disturbance regimes will therefore require understanding how the states and material legacies created by past disturbances condition ecosystem responses to those that follow.

## Materials and Methods

### Study Site and Permitting

All fieldwork was conducted on the north shore of Moorea, French Polynesia (17.54° S, 149.83° W), a small (∼ 16 km wide), triangular volcanic high island encircled by a barrier reef 1–1.5 km offshore that encloses a shallow lagoon. The Moorea Coral Reef Long Term Ecological Research (MCR LTER) project has collected time-series data on the physical environment and coral reef communities since 2005. Before the disturbances documented by the MCR LTER, the last major disturbance to substantially affect coral cover on Moorea was a severe bleaching event in 1991 (56). From 2007–2010, corals on the fore reef experienced a major outbreak of crown-of-thorns seastars (*Acanthaster planci*), followed immediately by Category 4 Cyclone Oli in February 2010, which scoured the north-shore fore reef to primary substrate (36). Thermal stress during a prolonged marine heat wave in the first half of 2019 caused mass coral mortality across the fore reef to at least 10 m depth (57), as well as sporadic coral mortality on patch reefs within the lagoon.

All experiments and other methods were performed in accordance with relevant guidelines and regulations for both French Polynesia and the University of California. Permits for field work were issued annually by the Haut-Commissariat de la République en Polynésie Française (DRRT) (Protocole d’Accueil, 2010-2026 to RJS and SJH).

### Effect of a Dead Skeleton Ghost Forest on Hysteresis in the Herbivore-Seaweed Relationship

Following the 2019 heat wave, we repeated and expanded a hysteresis experiment (test for path dependence) conducted at the same north-shore fore reef site after Cyclone Oli had denuded the reef (Fig. 1D) (29). The basic design imposed a gradient in herbivorous fish access to standardized substrate units using semipermeable exclusion cages. Cages were constructed from plastic-coated 2.5-cm wire mesh and measured 37 × 37 × 22 cm, with a benthic footprint of 0.14 m². Five herbivory treatments were created using cage openings of 2.5 × 2.5, 5 × 5, 7.5 × 7.5, or 10 × 10 cm and a partial-cage control (two sides and a partial top) that provided unfettered access by herbivores. We omitted a completely open treatment to quantify ambient herbivore pressure because previous experiments using this design detected no cage artifacts relative to completely open plots (29, 34); as a result, cage structure was present in all five herbivore treatments. Each replicate contained a standardized substrate consisting of four abutting, unglazed terra-cotta tiles (15.25 × 15.25 cm each) mounted with the rough undersides facing upward. Previous experiments demonstrated that these cage treatments generate a graded series of herbivorous fish visitation rates, bite rates, and biomass-weighted herbivory (31, 34). For visualization, treatments were labeled in descending order of herbivore activity as Ambient, 10, 7.5, 5, and None, where numbers indicate the maximum cage-opening dimension in centimeters.

To test for path dependence (hysteresis), we imposed the same herbivory gradient on two contrasting initial algal states: a thin layer of closely cropped turf algae characteristic of newly opened reef substrate and an established seaweed community composed primarily of the brown alga *Turbinaria ornata* (34). This design tested whether the level of herbivory required to prevent seaweed establishment differed from that required to remove established seaweeds, thereby estimating forward and reverse trajectories of the herbivore–seaweed relationship. For the *Turbinaria* initial state, small pieces of natural reef substrate bearing several plants were collected and attached underwater to a designated tile using nontoxic marine epoxy (Z-Spar A-788 Splash Zone Epoxy). Replicates were provisioned with approximately equal initial *Turbinaria* cover and biomass and then haphazardly assigned among herbivory treatments.

The post-Cyclone Oli experiment included only flat, structure-free patches, reflecting the scoured reefscape from which coral skeletons and rubble had been removed. We repeated this No-skeleton Patch treatment after the 2019 heat wave, but these experimental patches were now embedded within a reefscape containing an extensive ghost forest of standing dead coral skeletons (i.e., Heat Wave No-skeleton Patch treatment). We also added a Heat Wave Skeleton Patch treatment to test whether skeletal structure altered hysteresis locally. Each Heat Wave Skeleton Patch contained eight dead *Pocillopora* skeletons affixed to the tiles with epoxy. Skeletons had been cleaned, bleached, and dried to remove residual tissue and organic matter, ranged from 10–15 cm in height, and were arranged with small gaps between adjacent skeletons to mimic the natural spatial configuration of the surrounding ghost forest (12). Both patch-structure treatments received the same two initial algal states and five herbivory treatments, allowing us to test whether local skeletal structure altered the forward and reverse trajectories of the herbivore–seaweed relationship within the broader post-heat-wave ghost forest.

The fully factorial design crossed two within-patch structure treatments (No Skeleton and Skeleton), two initial algal states (Turf and *Turbinaria*), and five herbivory treatments, with six replicates per combination (2 × 2 × 5 × 6 = 120 cages). Replicates were deployed at the fore reef study site over six days in July 2022. Cages were continuously submerged in seawater during transport by boat to the fore reef and were affixed to the reef by divers using (removable) stainless-steel threaded rods and eyebolts embedded in the substratum. Replicates were haphazardly interspersed along the 10-12 m isobaths at the exact location of the post-cyclone hysteresis experiment (Fig. 1D) (29). Once deployed, cages were inspected periodically and cleaned of fouling organisms as needed. No cages were dislodged or damaged during the experiment. Seaweeds recruited and grew rapidly in Turf replicates exposed to the lowest level of herbivory. We used the same *a priori* criterion for terminating the experiment as in the previous hysteresis experiment following Cyclone Oli (29): the experiment ended when seaweed cover in the None × Turf treatment approached the initial cover in the *Turbinaria* treatment. This criterion was reached after 12 months, and the experiment was ended in July 2023. Cages were retrieved over a three-day period, transported to the laboratory in seawater, and placed in flowing seawater until they were processed within 24 hours of collection. Seaweed cover on the terra-cotta tiles was visually estimated by the same observer (SJH). All seaweeds were then removed by hand, sorted to genus, and weighed while damp (g) using a top-loading balance after excess water was removed using a standardized number of spins in a salad spinner.

For graphical presentation, seaweed cover and biomass were normalized separately within each initial state and patch-structure treatment by dividing each observation by the corresponding mean in the None herbivory treatment. To test for hysteresis, we determined whether the relationship between herbivory and final seaweed abundance depended on initial algal state. For each patch-structure treatment, we used permutational ANOVA to test the effects of Initial State, Herbivory, and their interaction on final seaweed cover and biomass. A small constant (0.01) was added to each percent cover value to accommodate zeros, after which cover was converted to a proportion and arcsine-square-root transformed prior to analysis. Permutational ANOVAs were implemented in R using the adonis2 function in the vegan package (58), with *P* values calculated from 999 permutations using unique sums of squares. We interpreted a significant Initial State × Herbivory interaction, accompanied by divergence between initial states at intermediate levels of herbivory, as evidence of hysteresis. To assess divergence between initial states, for each herbivory treatment below ambient we calculated the relative difference in final seaweed cover and biomass as the log ratio of the treatment means [log_10_(Seaweed) − log_10_(Turf)] and examined whether this difference increased (divergence) or decreased (convergence) with increasing herbivores (Fig. S2).

### Effect of Disturbance Type on State Transition Probabilities

We conducted two concurrent resilience experiments (tests for non-recovery) (33) on 60 lagoon patch reefs (Fig. 1D): a Coral Resilience Experiment that tested whether dead coral skeletons could facilitate a transition from a seaweed-depauperate to a seaweed-rich state, and a Seaweed Resilience Experiment that tested whether storm-like removal of established seaweeds could reverse the seaweed-rich state. Each experiment consisted of two treatments with 15 patch reefs per treatment. The site contains numerous small patch reefs (bommies; ∼ 1-4 m² surface area) situated 0.5-5 m apart (for more site details, see 29, 34). Replicates of the four treatment combinations were spatially interspersed throughout the study site.

For the Coral Resilience Experiment, we selected 30 patch reefs that lacked emergent seaweeds and were covered primarily by closely cropped turf and crustose coralline algae. The 30 reefs were haphazardly assigned to either the Coral Heat Wave or Coral Cyclone treatment (n = 15 each). The small amount of emergent biota initially present was removed from all reefs. To recreate the material legacy of coral mortality from a marine heat wave, we attached 10 dead branching *Pocillopora* skeletons to each designated Coral Heat Wave patch reef using marine epoxy (Z-Spar A-788 Splash Zone Epoxy) (Fig. 3B). Skeletons ranged from 9–15 cm in maximum diameter and 7–10 cm in height and were arranged to produce ∼ 50% cover and a distribution of gaps between adjacent skeletons that mirrored the distribution measured after the 2019 heat wave (12). Coral Cyclone reefs received no skeletons, representing the open reef condition produced by a scouring storm. After these initial manipulations, no further experimental interventions were made.

For the Seaweed Resilience Experiment, we selected 30 patch reefs with high cover (≥ 40%) of seaweeds, primarily *Turbinaria ornata* and *Sargassum pacificum*, that were similar in size, depth, and height above the seafloor. The 30 reefs were haphazardly assigned to either the Seaweed Cyclone or the Seaweed Control treatment (n = 15 each). Initial seaweed cover was compared between treatments using a Welch’s *t* test. To simulate storm-driven loss of established seaweeds, divers removed adults and juveniles by hand and then systematically removed holdfasts and small recruits with knives (Fig. 4B). After this initial manipulation, no further experimental interventions were made.

At the start of the experiments, we established a permanent 0.5 × 0.5 m sampling area at or near the top of each of the 60 patch reefs, matching the dimensions of the permanent quadrats used to quantify coral (59) and algal (60) cover on the fore reef in the MCR LTER long-term time series program. On Coral Heat Wave reefs, all experimentally added coral skeletons were positioned within this permanent sampling area (Fig. 3B). Divers periodically made visual estimates of seaweed cover within each sampling area and the fraction of experimentally added skeletons remaining. To classify reefs as seaweed-rich, we used the distribution of seaweed cover on 15 additional, unmanipulated reference reefs at the same site, where seaweeds had persisted as major space holders for approximately a decade (29, 34). Across surveys conducted during the experiment, the first quartile of seaweed cover on these reference reefs was 25.0% (Fig. S3), which we used as the threshold for classifying experimental reefs as seaweed-rich. Treatment differences in the probability of being seaweed-rich were evaluated separately for the two resilience experiments using Fisher’s exact tests.

Both resilience experiments were designed *a priori* to continue for at least four months after complete disappearance of the experimentally added *Pocillopora* skeletons. Four months approximates the turnover time of adult *T. ornata* in this system (29, 35), allowing us to test whether a seaweed-rich state established in the presence of skeletal refugia could persist through turnover of adult plants after those refugia disappeared. We quantified erosion of the experimentally added skeletons from photographs taken at 0, 13, 24, and 37 months and fitted a quadratic equation to the proportion of initially added skeletal material remaining through time. The model provided an excellent fit to the observed erosion trajectory (R² = 0.99) and predicted complete disappearance of the skeletons at approximately 42 months (Fig. S4). We therefore terminated both resilience experiments at 49 months, seven months after the predicted disappearance of the skeletal refugia and four months after visual confirmation of their complete disappearance. Thus, adult *T. ornata* had between one and nearly two turnover times after the skeletons were gone. The Coral Resilience Experiment was also sampled at 24 months to assess state transitions while substantial skeletal structure remained; slightly more than 60% of the initial skeletal material remained at that time.

### Natural Post-disturbance Transition Probabilities and Dynamics

The Moorea Coral Reef (MCR) Long Term Ecological Research (LTER) site has been documenting trends in the biota of Moorea’s fore reef since 2005. Sampling sites include two areas located approximately 1.5-2.0 km to the east and the west of our experimental fore reef site on the north shore (Fig. 1E). Diver surveys and photoquadrats in fixed, permanent quadrats (each 0.5 x 0.5 m) provide annual estimates of percent cover of major benthic substrate categories (e.g., coral, macroalgae, algal turf, sand), with most benthic taxa (including corals and seaweeds) resolved to species or genus level (59, 60). These data were used to create time series of cover of corals (59) and seaweeds (60) in the time periods following Cyclone Oli (2010) and the 2019 marine heat wave.

We quantified state-transition probabilities four years after each disturbance using the corresponding MCR LTER datasets. For coral, we used the Moorea Coral Reef LTER and Edmunds dataset (59), which contained 78 photoquadrats in 2014, four years after Cyclone Oli, and 73 (image acquisition failed for 5 quadrats) in 2023, four years after the 2019 marine heat wave. To establish an objective threshold for classifying quadrats as coral-rich, we used the same first-quartile convention applied to seaweeds. The threshold was calculated from the 2006 MCR LTER annual survey of six fore reef sites at 10 m depth, a time point when the fore reef community had not been subjected to a major disturbance for 15 years (56). The first quartile of coral cover in 2006 was 25.2% (median = 35.3%), which we used as the threshold for classifying quadrats as coral-rich. For seaweeds, we used the Moorea Coral Reef LTER and Carpenter dataset (60), which contained 100 quadrats in both 2014 and 2023. As in the resilience experiments, quadrats with ≥ 25.0% seaweed cover were classified as seaweed-rich. We then used Pearson’s chi-square tests with Yates’ continuity correction to test whether the probability of recovery to the coral-rich state and the probability of transition to the seaweed-rich state differed four years after Cyclone Oli versus four years after the 2019 marine heat wave.

Because coral and seaweed cover were estimated in both datasets we used (59, 60), we conducted an additional validation analysis to assess whether quadrats classified as coral-rich and seaweed-rich were effectively mutually exclusive. We pooled all 351 quadrats used in the four-year post-disturbance analyses (100 + 100 + 78 + 73) and determined how many simultaneously exceeded both the 25.2% coral-rich threshold and the 25.0% seaweed-rich threshold. The proportion of quadrats meeting both criteria provided an estimate of how frequently the two state classifications overlapped.

## Acknowledgments

We thank Thomas Adam, Deron Burkepile, Melissa Schmitt, Kelly Speare and Keenan Stears for constructive feedback and other contributions to this study, Hillary Krumbholz for assistance both with Information Management and extensive field support, and Lindsay Cullen, Madigan Boborci, Madeline Cunningham, Jordan Gallagher, Adam Goodman, Nick Jarymowycz, Anneke Padmos, Emalia Parlow, Katie Riley, Joaquin Sandoval, Marina Stoilova, and Xanthe Vitaz for their invaluable field and lab assistance, including long hours spent helping to construct, deploy, maintain, and retrieve field experimental infrastructure and processing samples. We also thank the staff of the University of California Gump Research Station and UC Enterprise Dive & Boat Safety Program for critical logistic support. This work is a contribution of the Moorea Coral Reef Long Term Ecological Research Site (MCR LTER) and was supported by National Science Foundation (grant OCE-2224354) as well as by research infrastructure provided by the Gordon and Betty Moore Foundation. Field research was completed under permits issued by the Territorial Government of French Polynesia (*Délégation à la Recherche*) and the *Haut-Commissariat de la République en Polynésie Française* (*Protocole d’Accueil* 2004-2026); we thank them for their continued support. With respect to the place name spelling of Moorea, we followed the *Raapoto* transcription system for *Reo M*ā*’ohi* (the traditional language of the Society Islands) that is adhered to by a large segment of the Tahitian community, but we also recognize other community members follow the *Te Fare Van*ā*’a* transcription system where the island name is spelled with an ’eta (*Mo’orea*) (for more background, see https://mcr.lternet.edu/about/mcr-site-policies/spelling-moorea).

## Author Contributions

Project co-leads (alphabetical): S.J.H., K.L.K., R.J.S.; Conceptualization and Research Design: All authors; Research Tasks: All authors; Data Analysis: A.J.B., K.L.K.; Visualizations: K.L.K., R.J.S.; Funding Procurement and Project Administration: S.J.H., R.J.S.; Writing: Initial draft: R.J.S.; Revisions: A.J.B., S.J.H., K.L.K.

## References

1. D. C. Reed et al., Responses of coastal ecosystems to climate change: Insights from long-term ecological research. BioScience 72, 871–888 (2022).

2. T. P. Hughes et al., Global warming and recurrent mass bleaching of corals. Nature 543, 373–377 (2017).

3. M. G. Turner, R. Seidl, Novel disturbance regimes and ecological responses. Annu. Rev. Ecol. Evol. Syst. 54, 63–83 (2023).

4. M. G. Turner, Disturbance and landscape dynamics in a changing world. Ecology 91, 2833–2849 (2010).

5. M. G. Turner et al., Climate change, ecosystems and abrupt change: Science priorities. Philos. Trans. R. Soc. B Biol. Sci. 375, 20190105 (2020).

6. T. P. Hughes et al., Global warming transforms coral reef assemblages. Nature 556, 492–496 (2018).

7. J. F. Johnstone et al., Changing disturbance regimes, ecological memory, and forest resilience. Front. Ecol. Environ. 14, 369–378 (2016).

8. T. P. Hughes et al., Ecological memory modifies the cumulative impact of recurrent climate extremes. *Nat*. Clim. Change 9, 40–43 (2019).

9. K. L. Kopecky et al., Legacies of foundation species shape life after death. Sci. Adv. 12, eaef9983 (2026).

10. K. Jõgiste et al., Hemiboreal forest: Natural disturbances and the importance of ecosystem legacies to management. Ecosphere 8, e01706 (2017).

11. K. L. Kopecky, A. C. Stier, R. J. Schmitt, S. J. Holbrook, H. V. Moeller, Material legacies can degrade resilience: Structure-retaining disturbances promote regime shifts on coral reefs. Ecology 104, e4006 (2023).

12. K. L. Kopecky, S. J. Holbrook, E. Partlow, M. Cunningham, R. J. Schmitt, Changing disturbance regimes, material legacies, and stabilizing feedbacks: Dead coral skeletons impair key recovery processes following coral bleaching. Glob. Change Biol. 30, e17504 (2024).

13. L. H. Gunderson, Ecological resilience—in theory and application. Annu. Rev. Ecol. Syst. 31, 425–439 (2000).

14. N. Knowlton, Thresholds and multiple stable states in coral reef community dynamics. Am. Zool. 32, 674–682 (1992).

15. D. R. Bellwood, T. P. Hughes, C. Folke, M. Nyström, Confronting the coral reef crisis. Nature 429, 827–833 (2004).

16. T. P. Hughes et al., Phase shifts, herbivory, and the resilience of coral reefs to climate change. Curr. Biol. 17, 360–365 (2007).

17. J. D. Woodley et al., Hurricane Allen’s impact on Jamaican coral reefs. Science 214, 749–755 (1981).

18. A. H. Sobel et al., Tropical cyclone frequency. Earth’s Future 9, e2021EF002275 (2021).

19. T. R. Knutson et al., Tropical cyclones and climate change assessment: Part II: Projected response to anthropogenic warming. Bull. Am. Meteorol. Soc. 101, E303–E322 (2020).

20. E. C. J. Oliver, et al., Marine heatwaves. Annu. Rev. Mar. Sci. 13, 313–342 (2021).

21. T. P. Hughes et al., Spatial and temporal patterns of mass bleaching of corals in the Anthropocene. Science 359, 80–83 (2018).

22. M. L. Harmelin-Vivien, The effects of storms and cyclones on coral reefs: A review. J. Coast. Res. 12, 211–231 (1994).

23. T. M. Kenyon et al., Coral rubble dynamics in the Anthropocene and implications for reef recovery. Limnol. Oceanogr. 68, 110–147 (2023).

24. D. E. Burkepile, M. H. Schmitt, K. Stears, M. K. Donovan, D. I. Thompson, Shared insights across the ecology of coral reefs and African savannas: Are parrotfish wet wildebeest? BioScience 70, 647–658 (2020).

25. K. L. Kopecky et al., Removing dead coral after marine heatwaves can mitigate coral–algae competition and increase viable coral recruitment. Ecol. Appl. 35, e70077 (2025).

26. S. J. Box, P. J. Mumby, Effect of macroalgal competition on growth and survival of juvenile Caribbean corals. Mar. Ecol. Prog. Ser. 342, 139–149 (2007).

27. D. B. Rasher, M. E. Hay, Chemically rich seaweeds poison corals when not controlled by herbivores. Proc. Natl. Acad. Sci. U.S.A. 107, 9683–9688 (2010).

28. P. J. Mumby, A. Hastings, H. J. Edwards, Thresholds and the resilience of Caribbean coral reefs. Nature 450, 98–101 (2007).

29. R. J. Schmitt, S. J. Holbrook, S. L. Davis, A. J. Brooks, T. C. Adam, Experimental support for alternative attractors on coral reefs. Proc. Natl. Acad. Sci. U.S.A. 116, 4372–4381 (2019).

30. M. Kayal et al., Predator crown-of-thorns starfish (*Acanthaster planci*) outbreak, mass mortality of corals, and cascading effects on reef fish and benthic communities. PLoS ONE 7, e47363 (2012).

31. S. J. Holbrook, R. J. Schmitt, T. C. Adam, A. J. Brooks, Coral reef resilience, tipping points and the strength of herbivory. Sci. Rep. 6, 35817 (2016).

32. S. J. Holbrook et al., Recruitment drives spatial variation in recovery rates of resilient coral reefs. Sci. Rep. 8, 7338 (2018).

33. V. Dakos, S. Kéfi, Ecological resilience: What to measure and how. Environ. Res. Lett. 17, 043003 (2022).

34. R. J. Schmitt, S. J. Holbrook, A. J. Brooks, T. C. Adam, Evaluating the precariousness of coral recovery when coral and macroalgae are alternative basins of attraction. Limnol. Oceanogr. 67, S285–S297 (2022).

35. S. L. Davis, “Mechanisms underlying macroalgal phase shifts in coral reef ecosystems,” PhD dissertation, University of California, Santa Barbara, CA (2016).

36. T. C. Adam et al., Herbivory, connectivity, and ecosystem resilience: Response of a coral reef to a large-scale perturbation. PLoS ONE 6, e23717 (2011).

37. T. C. Adam et al., Priority effects in coral–macroalgae interactions can drive alternate community paths in the absence of top-down control. Ecology 103, e3831 (2022).

38. G. Tsounis, P. J. Edmunds, The potential for self-seeding by the coral *Pocillopora* spp. in Moorea, French Polynesia. PeerJ 4, e2544 (2016).

39. T. C. Adam et al., How will coral reef fish communities respond to climate-driven disturbances? Insight from landscape-scale perturbations. Oecologia 176, 285–296 (2014).

40. X. Han, T. C. Adam, R. J. Schmitt, A. J. Brooks, S. J. Holbrook, Response of herbivore functional groups to sequential perturbations in Moorea, French Polynesia. Coral Reefs 35, 999–1009 (2016).

41. S. Bennett, A. Vergés, D. R. Bellwood, Branching coral as a macroalgal refuge in a marginal coral reef system. Coral Reefs 29, 471–480 (2010).

42. L. D. Puk, A. Marshell, J. Dwyer, N. R. Evensen, P. J. Mumby, Refuge-dependent herbivory controls a key macroalga on coral reefs. Coral Reefs 39, 953–965 (2020).

43. R. Seidl, W. Rammer, T. A. Spies, Disturbance legacies increase the resilience of forest ecosystem structure, composition, and functioning. Ecol. Appl. 24, 2063–2077 (2014).

44. S. L. Davis, Associational refuge facilitates phase shifts to macroalgae in a coral reef ecosystem. Ecosphere 9, e02272 (2018).

45. J. P. Gallagher, R. J. Schmitt, D. E. Burkepile, S. J. Holbrook, Nutrient enrichment strengthens feedbacks that promote and stabilize coral-to-macroalgae regime shifts on tropical reefs. Ecosphere 17, e70642 (2026).

46. C. J. Briggs, T. C. Adam, S. J. Holbrook, R. J. Schmitt, Macroalgae size refuge from herbivory promotes alternative stable states on coral reefs. PLoS ONE 13, e0202273 (2018).

47. D. T. Cook, S. J. Holbrook, R. J. Schmitt, Insights from the competition–palatability trade-off paradigm on the reversibility of coral-to-macroalgae regime shifts. Sci. Rep. (2026), doi:10.1038/s41598-026-59166-7.

48. M. Scheffer, S. Carpenter, J. A. Foley, C. Folke, B. Walker, Catastrophic shifts in ecosystems. Nature 413, 591–596 (2001).

49. M. Scheffer, S. R. Carpenter, Catastrophic regime shifts in ecosystems: Linking theory to observation. Trends Ecol. Evol. 18, 648–656 (2003).

50. K. R. N. Anthony et al., Operationalizing resilience for adaptive coral reef management under global environmental change. Glob. Change Biol. 21, 48–61 (2015).

51. Y.-M. Bozec, P. J. Mumby, Synergistic impacts of global warming on the resilience of coral reefs. Philos. Trans. R. Soc. B Biol. Sci. 370, 20130267 (2015).

52. A. K. Ford, M. Hamilton, Y. Nand, M. Puotinen, S. D. Jupiter, S. Dulunaqio, W. Naisilisili, S. Mangubhai, Comparing impacts and recovery of locally managed reefs after exposure to extreme waves from a category 5 cyclone. Coral Reefs 44, 1909–1926 (2025).

53. M. J. Emslie et al., Increasing disturbance frequency undermines coral reef recovery. Ecol. Monogr. 94, e1619 (2024).

54. A. K. Cresswell et al., Coral reef state influences resilience to acute climate-mediated disturbances. Glob. Ecol. Biogeogr. 33, 4–16 (2024).

55. J. S. Harding, E. F. Benfield, P. V. Bolstad, G. S. Helfman, E. B. D. Jones III, Stream biodiversity: The ghost of land use past. Proc. Natl. Acad. Sci. U.S.A. 95, 14843–14847 (1998).

56. M. L. Trapon, M. S. Pratchett, L. Penin, Comparative effects of different disturbances in coral reef habitats in Moorea, French Polynesia. J. Mar. Sci. 2011, 807625 (2011).

57. K. E. Speare, T. C. Adam, E. M. Winslow, H. S. Lenihan, D. E. Burkepile, Size-dependent mortality of corals during marine heatwave erodes recovery capacity of a coral reef. Glob. Change Biol. 28, 1342–1358 (2022).

58. J. Oksanen et al., *vegan: Community Ecology Package*. R package version 2.6–2 (2022).

59. Moorea Coral Reef LTER and P. Edmunds. 2026. MCR LTER: Coral Reef: Long-term Population and Community Dynamics: Corals, ongoing since 2005 ver 44. Environmental Data Initiative. 10.6073/pasta/65f9aad0f4332a144238593e28be395d.

60. Moorea Coral Reef LTER and R. Carpenter. 2025. *MCR LTER: Coral Reef: Long-term Population and Community Dynamics: Benthic Algae and Other Community Components, ongoing since* 2005 ver 38. Environmental Data Initiative. 10.6073/pasta/44d263de042710767ed770986fd12f5c.

