## Supporting information for "Heat Waves Promote, but Cyclones Reverse, Coral-to-Seaweed Regime Shifts by Reshaping Reef Resilience"

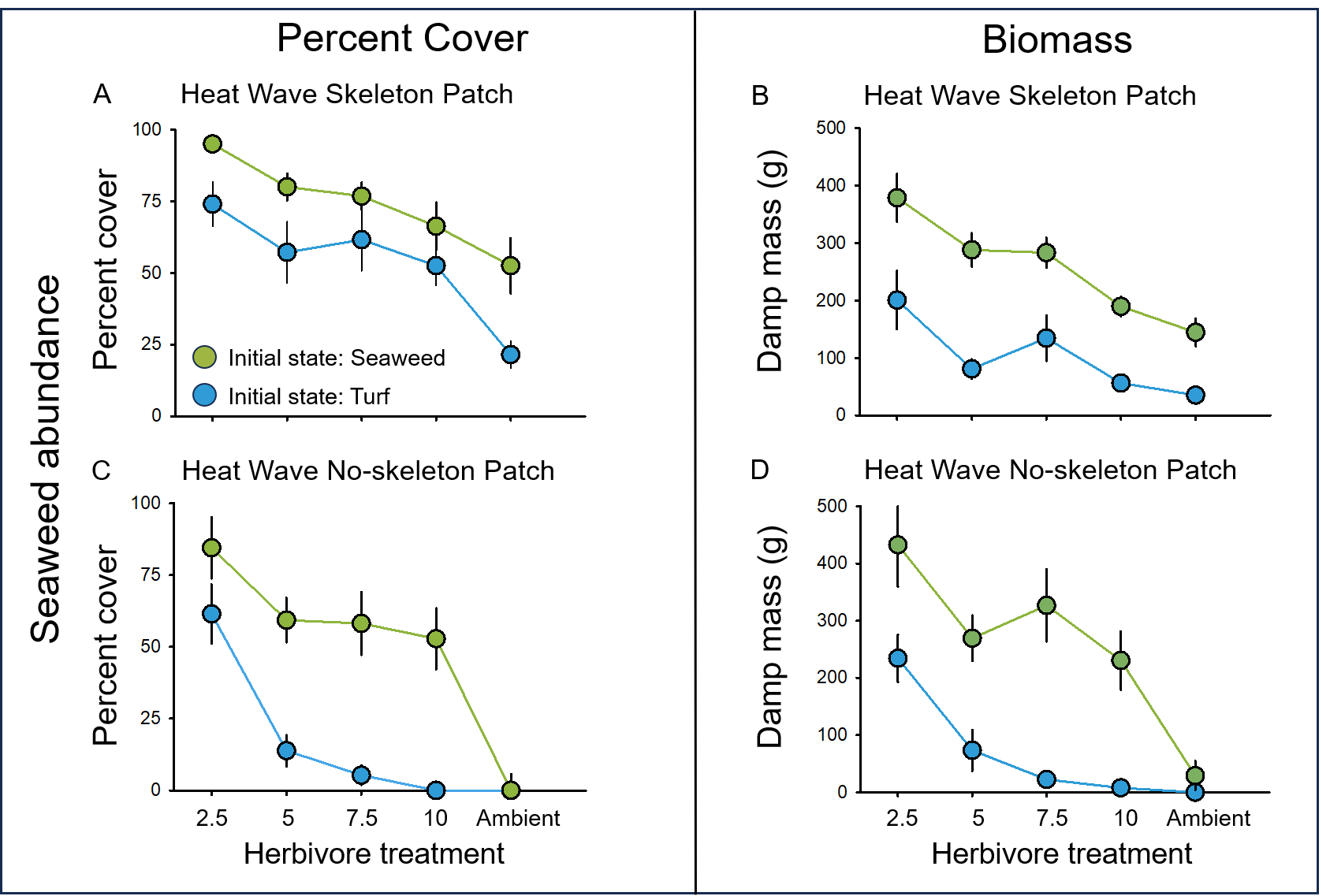

**Figure S1.** Results of the Heat Wave Skeleton Patch (A, B) and Heat Wave No-skeleton Patch (C, D) treatments of the Hysteresis Experiments, showing final seaweed cover (A, C) and damp biomass (B, D) across the experimental herbivore gradient as a function of initial algal state (Turf: blue circles; Seaweed: green circles). Data are means ± 1 SE (n = 6). All experimental patches were embedded within the post-heat-wave ghost forest of standing dead coral skeletons, composed primarily of *Pocillopora*, that resulted from mass bleaching mortality during the severe 2019 marine heat wave. In the Heat Wave Skeleton Patch treatments (A, B), physical refugia were present both within an experimental patch and throughout the surrounding ghost forest; dead *Pocillopora* skeletons in each replicate matched the density and spacing of those in the surrounding reefscape. In the Heat Wave No-skeleton Patch treatments (C, D), replicates lacked physical refugia from dead coral skeletons but were embedded within the surrounding ghost forest. See Table S1 for results of permutational ANOVAs testing the effects of herbivory treatment, initial algal state, and their interaction. See Fig. S2 for evidence that the significant interaction in the Heat Wave No-skeleton Patch treatments resulted from divergence between initial algal states as herbivore abundance increased.

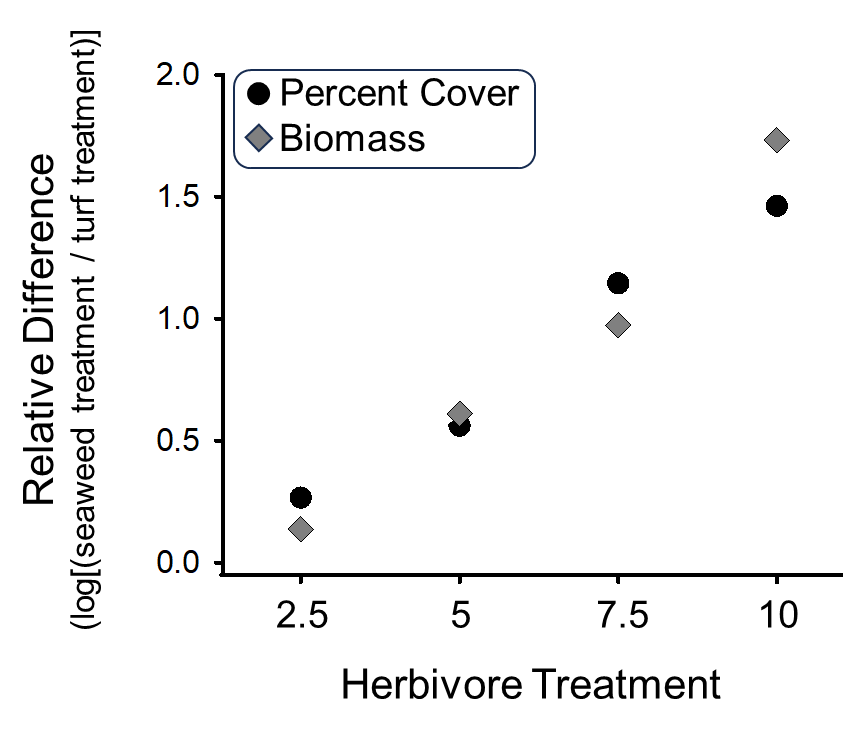

**Figure S2.** Divergence between initial algal states in the Heat Wave No-skeleton Patch Experiment. For both measures of final seaweed abundance (percent cover: black circles; damp biomass: gray diamonds), the significant interaction between initial algal state (Turf vs. Seaweed) and herbivory treatment resulted from increasing divergence between initial states as herbivore abundance increased. Shown are the relative differences in final seaweed abundance between initial states, calculated for each herbivory treatment below ambient as log ratios of the treatment means [log_10_(Seaweed) − log_10_(Turf)], using non-normalized data. Positive values indicate greater final seaweed abundance in the Seaweed than Turf initial state, and the positive relationship with increasing herbivore abundance indicates that final seaweed abundance increasingly depended on initial algal state, consistent with hysteresis and a region of coral–seaweed bistability.

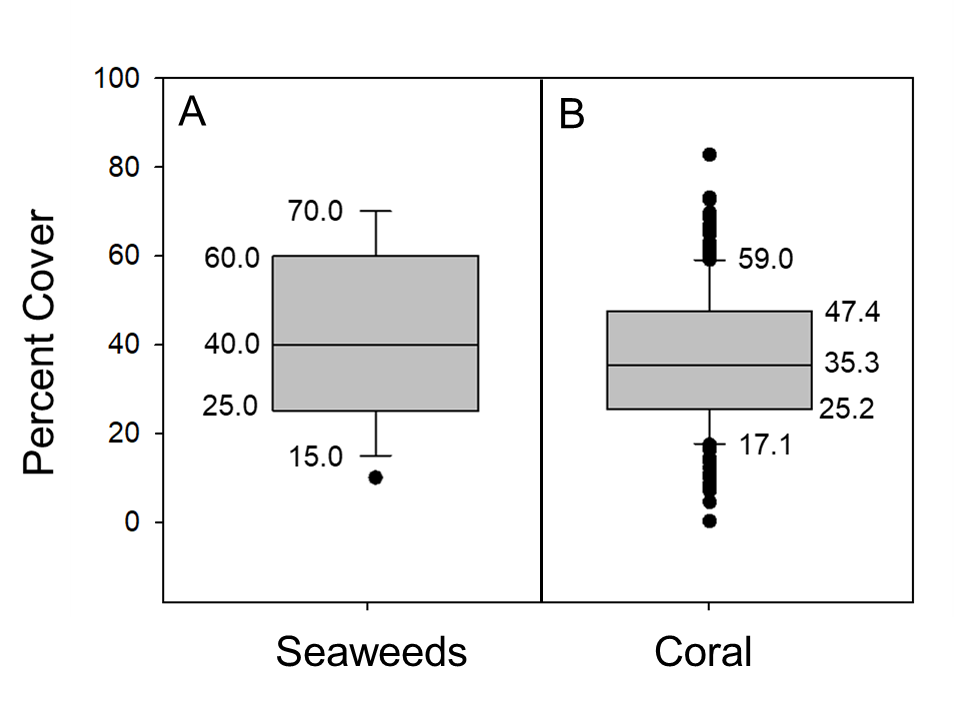

**Figure S3.** Reference distributions used to establish thresholds for classifying experimental and long-term survey plots as seaweed-rich or coral-rich. (A) Distribution of seaweed cover in 2022 on 15 unmanipulated reference patch reefs at the lagoon study site that had remained seaweed-rich for approximately a decade. The box plot shows the lower extreme, first quartile, median, third quartile, and upper extreme; points denote outliers. The first quartile (25.0% seaweed cover) was used as the threshold for distinguishing seaweed-depauperate from seaweed-rich states. (B) Distribution of live coral cover at 10 m depth on the fore reef of Moorea in 2006, following 15 years without a major disturbance to the fore reef coral community. The first quartile (25.2% live coral cover) was used as the threshold for distinguishing coral-depauperate from coral-rich states. Data are from the MCR LTER core time-series program (59, 60) (Moorea Coral Reef LTER and P. Edmunds. 2026. MCR LTER: Coral Reef: Long-term Population and Community Dynamics: Corals, ongoing since 2005, ver. 44. Environmental Data Initiative (<https://doi.org/10.6073/pasta/65f9aad0f4332a144238593e28be395d>).

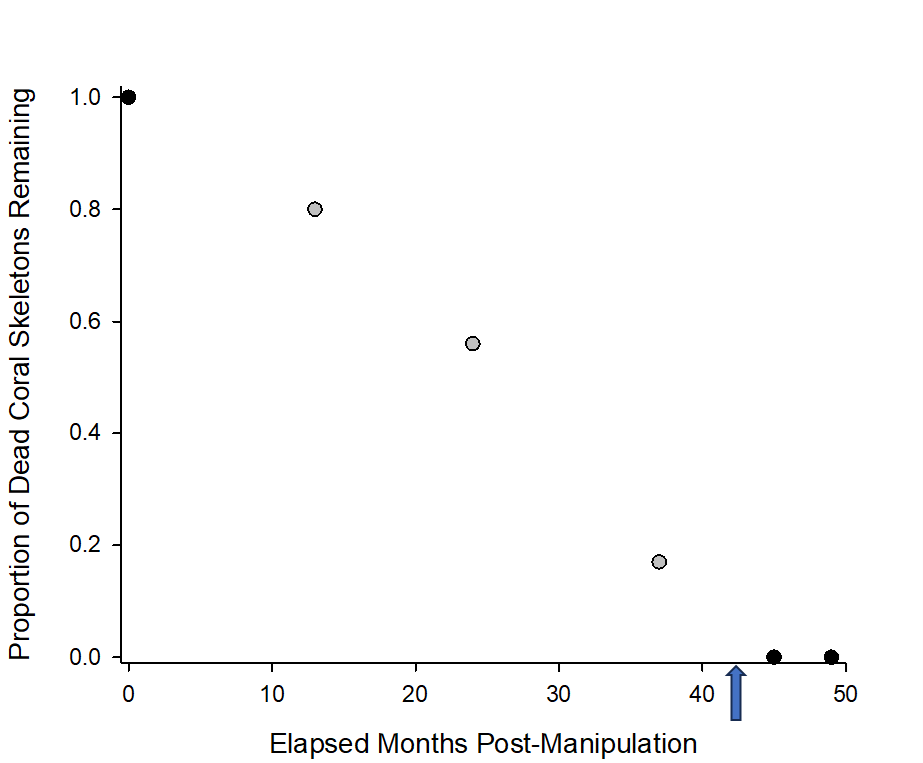

**Figure S4.** Erosion of experimentally added dead *Pocillopora* skeletons on Coral Heat Wave plots in the Coral Resilience Experiment. Points show estimates of the proportion of initially added skeletal material that remained, derived from photographs taken at 0, 13, 24, and 37 months after the start of the experiment. A quadratic model fitted to these four estimates (R² = 0.99) predicted complete disappearance of skeletal material at approximately 42 months (blue arrow). Complete disappearance of the experimental skeletons was confirmed at 45 months, and the experiment was terminated at 49 months.

**A. Heat Wave Skeleton Patch**

Seaweed Percent Cover	Seaweed Biomass

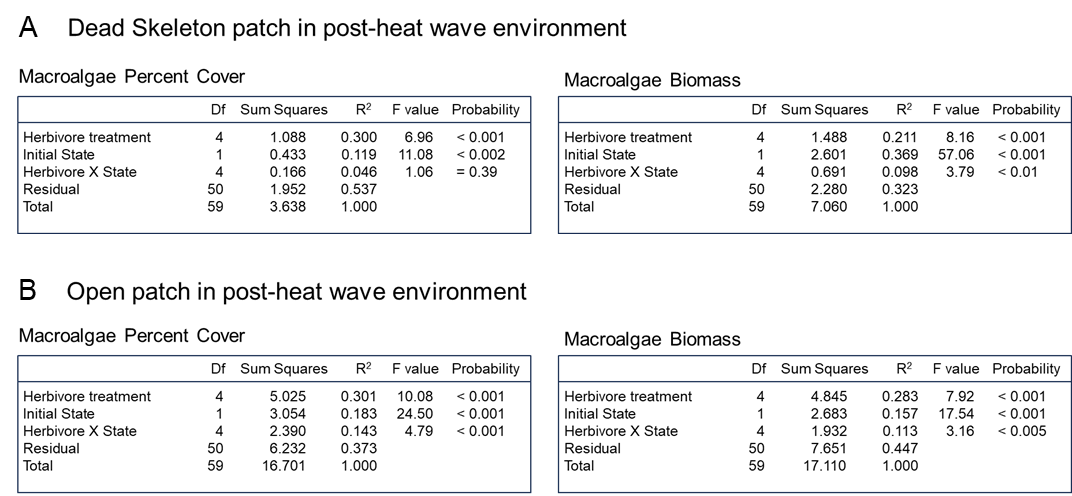

**B. Heat Wave No-skeleton Patch**

Seaweed Percent Cover	Seaweed Biomass

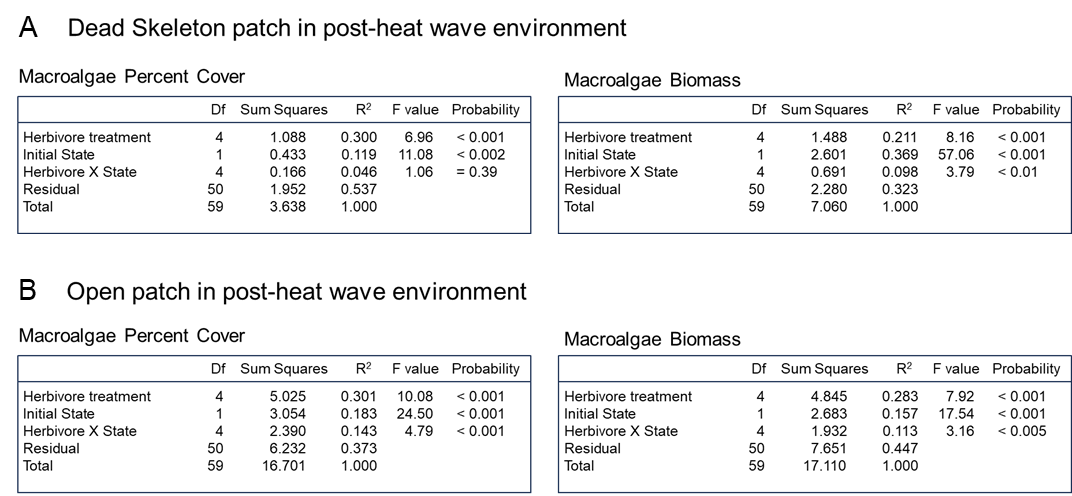

**Table S1.** Results of Permutational ANOVAs for the Hysteresis Experiment on the fore reef that tested for effects of herbivory treatment and initial community state, and for an interaction between the two factors. Separate models were run for open patches and patches with dead skeletons, as well as for seaweed percent cover and biomass as response variables.
